# Cross-species analysis between the maize smut fungi *Ustilago maydis* and *Sporisorium reilianum* highlights the role of transcriptional plasticity of effector orthologs for virulence and disease

**DOI:** 10.1101/2020.11.03.366443

**Authors:** Weiliang Zuo, Jasper RL Depotter, Deepak K Gupta, Marco Thines, Gunther Doehlemann

## Abstract

- The constitution and regulation of effector repertoires determines and shapes the outcome of the interaction with the host. *Ustilago maydis* and *Sporisorium reilianum* are two closely related smut fungi, which both infect maize, but cause distinct disease symptoms. Understanding how effector orthologs are regulated in these two pathogens can therefore provide insights to pathogen evolution and host adaption.
- We tracked the infection progress of *U. maydis* and *S. reilianum* in maize leaves, characterized two distinct infection stages for cross species RNA-sequencing analysis and identified 207 out of 335 one-to-one effector orthologs being differentially regulated during host colonization, while transcriptional plasticity of the effector orthologs correlated with the distinct disease development strategies.
- By using CRISPR-Cas9 mediated gene conversion, we identified two differentially expressed effector orthologs with conserved function between two pathogens. Thus, differential expression of functionally conserved genes contributes to species specific adaptation and symptom development. Interestingly, another differentially expressed orthogroup (*UMAG_05318/Sr1007*) showed diverged protein function during speciation, providing a possible case for neofunctionalization.
- Collectively, we showed the diversification of effector genes in related pathogens can be caused both by plasticity on the transcriptional level, as well as through functional diversification of the encoded effector proteins.

## Introduction

During symbiosis, microbes secret effector proteins to facilitate a compatible interaction with their hosts. Effectors hold various functions, such as the suppression of host immunity and host metabolism manipulation to promote host infection. Co-evolution of pathogens with their hosts is driven by an arms race causing selection pressure on perceived effector genes against their recognition by the host immune system (Jones & Dangl, 2006).

Polymorphisms of effectors are diverse, including presence/absence variation, amino acid substitution, epigenetic modification and transcriptional plasticity, which together determine the outcome of the plant-pathogen interaction (Gijzen *et al*., 2014; Toruño *et al*., 2016; Franceschetti *et al*., 2017; Torres *et al*., 2020). Until now, many effector studies have focused on genome sequencing and effector prediction and variation amongst phylogenetically related pathogens (Raffaele & Kamoun, 2012; Sánchez-Vallet *et al*., 2018); functional elucidation of single effector proteins (Selin *et al*., 2016); as well as transcriptional changes during host infection in one-to-one host-pathogen interactions (Lanver *et al*., 2018). However, information available on the transcriptional regulation of effector orthologs across related pathogen species is scarce.

Smut fungi are a group of fungal plant pathogens consisting of more than 1,500 species, which infect mainly grasses including major cereals such as maize, sorghum, wheat, barley and sugar cane (Begerow *et al*., 2004). A characteristic feature of many smut fungi is their dimorphic growth, where distinct growth states are observed: saprophytic growth as yeast, and filamentous hyphae which initiate the biotrophic interaction with the host (Kämper *et al*., 2006a). Smuts often infect their host through the root or coleoptile (Martinez *et al*., 2002; Laurie *et al*., 2012), from where the fungi proliferate without evident symptoms during the early vegetative growth stage of the host. When plants switch into the reproductive phase, fungal hyphae residing in floral structure may produce massive black/ brown teliospores (the notorious “smut” phenotype). This type of infection style is also found for the smut fungus *Sporisorium reilianum*, which infects either sorghum (*S.□reilianum f. sp. reilianum*) or maize (*S.□reilianum f. sp. zeae*) and causes head smut disease. Closely related to this is *Ustilago maydis*, which causes common maize smut disease (Kämper *et al*., 2006a; Schirawski *et al*., 2010). While *U. maydis* has the ability to cause disease symptoms on all the aerial maize tissues, *S.□reilianum* spreads systemically in the plant while it stays close to the vascular bundles and grows up the main stem axis to reach the cob primordia. *Ustilago. maydis* causes locally restricted plant tumors within a short time of less than 2 weeks under laboratory conditions. Comparative genome analysis of *U. maydis* with *S. reilianum* revealed that the two maize smuts share high genome similarity with regard to gene number and synteny (Schirawski *et al*., 2010). Furthermore, previous studies suggested that, similar to *U. maydis, S. reilianum* can efficiently infect maize leaves to spread systemically and cause head smut disease in floral organs. In recent years, several virulence effectors were identified using seedling infection approaches (Schirawski *et al*., 2010; Ghareeb *et al*., 2015; Schweizer *et al*., 2018). Together, this renders the two closely related species a well-suited model to study regulation and function of orthologous genes.

In *U. maydis*, the expression of effector genes is precisely regulated via a network of hierarchical transcriptional factors (TF), which modulates activity of the transcription of effectors genes at distinct stages of infection (Skibbe *et al*., 2010; Lanver *et al*., 2018). The *b* mating type locus encoded genes, *bE/bW*, form a heterodimer TF, which not only regulates the expression of 38 effectors, but also trigger the expression of several TFs including a C2H2 zinc finger TF Rbf1 (regulator of b-filament) (Heimel *et al*., 2010b,a), which then turn on the TF Hdp2 (homeodomain transcription factor 2) and Biz1 (*b*-dependent zinc finger protein), together activating the expression of effectors important for initialing the biotrophic interaction (Flor-Parra *et al*., 2006; Heimel *et al*., 2010b). At later infection stages, the forkhead TF Fox1 controls effectors necessary for full virulence and inhibition of the host defense after establishment of biotrophic interaction (Zahiri *et al*., 2010) and the WOPR TF Ros1 (Tollot *et al*., 2016) regulate effector gene expression related to fungal sporogenesis in mature tumors. A comprehensive set of RNA-seq data covering the whole biotrophic growth phase of *U. maydis* identified distinct expression patterns of effectors and a novel virulence related TF Nlt1 (no leaf tumors1) (Lanver *et al*., 2018). The expression of effectors in *U. maydis* was also found to be regulated in an organ-specific manner (Skibbe *et al*., 2010), and a more recent study using RNA-sequencing from laser capture micro-dissected cells revealed a host cell-type specific expression pattern for several effector genes (Matei *et al*., 2018).

Reverse genetics studies identified various *U. maydis* effectors holding distinct virulence functions, and some of these effectors have been functionally characterized on the molecular level (Lanver *et al*., 2017). Examples are the apoplastic effectors Pit2 and Pep1, which inhibit the maize apoplastic Papain-like cysteine proteases (PLCPs) (Mueller *et al*., 2013; Misas Villamil *et al*., 2019) and peroxidase activities (Hemetsberger *et al*., 2012), respectively. The effectors Cmu1 (Djamei *et al*., 2011) and Tin2 (Tanaka *et al*., 2014) are translocated into the host cytoplasm to manipulate host SA and lignin metabolism, respectively. The leaf-specific effector See1 re-activates host DNA synthesis to directly promote tumor formation in bundle sheath cells (Redkar *et al*., 2015a; Matei *et al*., 2018). In addition, the secreted repetitive effector protein Rsp3 coats the fungal cell wall and protects it against a host anti-fungal protein (Ma *et al*., 2018). In contrast, only one *S. reilianum* specific effector SAD1suppresses the apical dominance of infected maize to increase branch number has been functional characterized in *S. reilianum* (Ghareeb *et al*., 2015), and several species-specific and cluster 19A effectors were identified to contribute virulence (Ghareeb *et al*., 2019). While these studies focused on the effectors’ function on the protein level, it remains unknown if and how transcriptional regulation of effectors contributes to virulence and infection style of the pathogen.

Comparative genomic analysis from phylogenetically related plant pathogens suggesting the effector repertories of smut fungi can be classified as core and accessory effector based on the conservation level (Schuster *et al*., 2018; Beckerson *et al*., 2019; Depotter & Doehlemann, 2020; Depotter *et al*., 2020). Core effectors are conserved in all phylogenetically related pathogen and may be involved in conserved biological process vital for infection, leading to stabilization of such effectors during speciation (Hemetsberger *et al*., 2015; Irieda *et al*., 2019; Thines, 2019). Complementary, accessory effectors are less conserved and only found in few or individual pathogen species and these were supposed to have more subtle/specific functions in virulence. Previous studies suggested that effector orthologs between *U. maydis* and *S. reilianum* are more functionally conserved compared to *Ustilago hordei* and/or *Melanopsichium pennsylvanicum* (Sharma *et al*., 2014; Redkar *et al*., 2015b; Stirnberg & Djamei, 2016).

Clustered regularly interspaced short palindromic repeats (CRISPR)-Cas9 was first identified in *Streptococcus pyogenes* as part of the bacterial immune system against bacteriophage infection (Barrangou *et al*., 2007), and has been widely adapted as genome editing tool in various organisms including filamentous fungi and oomycetes (Schuster & Kahmann, 2019). The Cas9 endonuclease generates DNA double-strand break in a specific target that is recognized by the sgRNA (single guide RNA). This double strand break can be repaired by non-homologous end-joining (NHEJ) which eventually results in a gene knock-out mutant. Alternatively, homology direct repair (HDR) with a donor template can be used for the generation of knock-in or gene conversion mutants. In *U. maydis*, the CRISPR-Cas9 is achieved by expressing the codon-optimized Cas9 protein and sgRNA in an autonomous replication plasmid (Schuster *et al*., 2016, 2018), which shows high efficiency to generate a selection marker free genome modification mutant. The off-target effects are further reduced by application of high fidelity variant Cas9HF1 (Zuo *et al*., 2020).

In this study, we examined the growth of *U. maydis* and *S. reilianum* during leaf infection for recording similar and divergent biotrophic interaction with maize. We investigated the regulation of orthologous genes with an emphasis on effector genes.

## Materials and Methods

### Strains, growth conditions and plant infection

The mating compatible isolates of *U. maydis* FB1, FB2 and *S. reilianum* SRZ1, SRZ2 were used for RNA-seq, and the solo-pathogenic *U. maydis* strain SG200 and its respective knock-out and ortholog conversion mutants were used for virulence tests. All strains were grown in YEPS light liquid medium with 200 rpm shaking or potato dextrose agar (PD, Difco) plate at 28°C. *Escherichia coli* strain Top10 was used for cloning purpose and grown in dYT liquid medium or YT agar plate with supplement of corresponding antibiotics. Maize variety Early Golden Bantam was used for infection. The plants were grown in a controlled condition of 16 h light at 28 °C and 8 h dark at 22°C. 7 days-old seedlings were infected with a mixture of compatible *U. maydis/S. reilianum* isolates or SG200, and disease symptoms were scored at 12 days after infection.

### Staining and microscopy

For microscopy purposes, 0.1% tween-20 was added in the inoculum. To visualize the appressorium, leaves were stained 20 hours post infection with calcofluor solution (100 μg/ml) for 1 min and briefly rinsed with water (Lanver *et al*., 2014). To visualize the fungal growth inside the infected leaves, WGA-AF488 (Wheat Germ Agglutinin, Alexa Fluor 488) and Propidium Iodide were used for staining the fungal and plant cell wall, respectively as previously described (Doehlemann *et al*., 2009a). The microscopy was done on a Nikon Eclipse Ti Inverted Microscope with the Nikon NIS-ELEMENTS software (Düsseldorf, Germany) and the photos were taken by HAMAMATSU camera. Confocal microscopy was done by Leica TCS SP8 confocal laser scanning microscope (Leica, Wetzlar, Germany) and the following filters were used excitation 458 nm and emission 470-490 nm for WGA-AF488 and excitation 561 nm and emission 590-603 nm for Propidium Iodide.

### RNA preparation and RNA-seq

For RNA preparation, 2-cm long of 3^rd^ leaf sections from more than 15 individual plants were collected for each sample. The compatible haploid *U. maydis* and *S. reilianum* cells from cultures with OD_600_ around 0.8 were spun down and mixed in a 1:1 ratio as axenic culture (AC) control. The plant tissues and cell pellets were ground into fine powder with liquid nitrogen and the RNAs were prepared using TRizol (Thermo Fisher, Waltham, USA) according to the manufacture’s protocol and followed by DnaseI digestion (Thermo Fisher, Waltham, USA). The RNA libraries were prepared using an Illumina TruSeq Stranded mRNA kit (Illumina, San Diego, USA), and paired-end sequencing was performed on the a HiSeq4000 platform to produce 2×75 bp long reads at the Cologne Center for Genomics (Cologne, Germany). Three independent biological replicates have been sampled and used for RNA-seq analysis.

### Data analysis

For the gene expression analysis of *U. maydis* and *S. reilianum* individually, reads of three biological replicates were filtered using the Trinity software (v2.9.1) option trimmomatic under the standard settings (Grabherr *et al*., 2013). They were mapped to a reference assembly using Bowtie 2 (v2.3.5.1) with the first 15 nucleotides on the 5’-end of the reads being trimmed (Langmead & Salzberg, 2012). The reference genome was either the genome assembly of *U. maydis* (Kämper *et al*., 2006b) or *S. reilianum* (Schirawski *et al*., 2010) combined with that of *Z. mays* B73 version 3 (Schnable *et al*., 2009). Reads were counted to the *U. maydis* and *Z. mays* loci using the R package Rsubread (v1.34.7) (Liao *et al*., 2019). edgeR package v3.26.8 was used for statistical analysis of differential gene expression and pairwise comparison was conducted using Generalized linear models (glm) (Robinson *et al*., 2009). Genes with log2 fold change >1 and *p*<0.05 were considered as differentially regulated between timepoints, and genes with log2 fold change <1, *p*<0.05 or log2 fold change >1 but *p*>0.05 were considered as not significantly differenr in the induced pattern analysis. The induced patterns of effectors were defined based on the differentially regulation relationships between AC, 2 dpi and 4 dpi samples with following criterias: pattern1 are genes induced *in planta* (expression levels at 2 dpi and 4 dpi were significantly higher than AC samples) but the 4 dpi expression level is higher than 2 dpi; genes classified as pattern 2 were induced *in planta*, but the expression levels were similar between 2 and 4 dpi; pattern 3 genes were only induced at 2 dpi and had dropped again at 4 dpi to AC level; pattern 4 genes were significantly induced *in planta* at both timepoints, but the expression level of 2 dpi is higher than 4 dpi; pattern 5 genes are only induced at 4 dpi.

A phylogenetic tree was constructed including the smut species *U. hordei* (Laurie *et al*., 2012), *M. pennsylvanicum* (Sharma *et al*., 2014), *S. scitamineum* (Taniguti *et al*., 2015), *S. reilianum* (Schirawski *et al*., 2010) and *U. maydis* (Kämper *et al*., 2006a). *Moesziomyces antarcticus* was used as an outgroup (Morita *et al*., 2013). A phylogenetic tree was constructed based on 1643 Benchmarking Universal Single-Copy Orthologs (BUSCOs) from the database “basidiomycota_odb10” that were present in single copy in all members investigated in the phylogenetic reconstruction (Seppey *et al*., 2019). For every gene, orthologs were aligned using MAFFT (v7.464) option “--auto” (Katoh & Standley, 2013). These aligned gene sequences were then concatenated for every species and used for tree construction using RAxML (v8.2.11) with the substitution model “GTRGAMMA” and 100 bootstraps (Stamatakis, 2014).

For comparison of ortholog expression, high quality paired end transcriptomic reads were mapped using Bowtie2 with default parameters over all orthologous genes of *Ustilago maydis* and *Sporisorium reilianum* as calculated using OrthoMCL. Average coverage of mapping of each gene was calculated from the sorted BAM file by summing the coverage of each base of a gene and then dividing this by the length of that gene. The relative expression value of a gene was calculated by using the following formula: 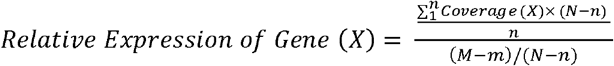, Where n= Length of the gene (X), N = Total number of bases in all genes; m= Number of bases mapped to gene (X); M= Number of bases mapped to all genes. *Student t-test* and/or one sample *t-test* were used for significance test with Benjamini-Hochberg *p* value correction for multiple comparison. Orthologs with fold change >3 and adjust *p*<0.05 were considered as differentially regulated ortholog between *U. maydis* and *S. reilianum*.

### Gene ontology and GO enrichment analysis

The gene ontology classification and overrepresentation analysis were done by using the *U. maydis* annotation data from PANTHER (pantherdb.org). The overrepresentation was conducted by using *Fisher’s exact test* with Bonferroni correction.

### Quantitative Real-Time PCR

For biomass quantification, the DNA of infected leaves from three biological replicates was prepared by Buffer A (0.1M Tris-HCl, 0.05M EDTA, 0.5M NaCl, 1.5% SDS), then further purified by MasterPure Complete DNA and RNA Purification Kit Bulk Reagents (Epicentre, Madison, USA). 100 ng of DNA was used for qPCR, and 2^-ΔCt^ was calculated to determine the ratio between fungal *peptidylprolyl isomerase* (*ppi*) and maize *GAPDH*.

For gene expression, the total RNA was reverse-transcribed with Oligo(dT) primer by using RevertAid First Strand cDNA Synthesis Kit (Thermo Scientific, Waltham, USA), and the expression level of each effectors were determined by 2^-ΔCt^ ratio between effector genes and *ppi* genes from respective species. The qPCR was conducted in CFX96 Real-Time PCR Detection System (Bio-Rad, Hercules, USA) with GoTaq qPCR mix (Promega, Madison, USA). The primers designed in this study were listed in Table. **S1**.

### Gene conversion in *U. maydis* by CRISPR and disease scoring

CRISPR-Cas9 mediated genome editing was used to generate the effector ortholog conversion mutants in *U. maydis* solo-pathogenic strain SG200. An sgRNA was designed to target the coding region or the promoter region of the effector to generate the double strand break, which was then repaired by homology-directed repair using a donor plasmid as template. The design of sgRNA and cloning was done as previously described (Zuo *et al*., 2020). To construct donor plasmid, 1 kb fragments flanking the effector coding region or promoter regions and the respective *S. reilianum* ortholog open reading frame or promoters were amplified by Phusion DNA polymerase (NEB, Ipswich, USA) with primers (Sigma Aldrich, St. Louis, US) listed in Table. **S1** and cloned into pAGM1311 vector by Gibson assembly (New England Biolabs, Ipswich, USA). The CRISPR plasmid and circular donor plasmid were co-transformed into SG200 protoplast and mutants were singled out on PD plate with 2 μg/ml carboxin for CRISPR plasmid first then transfer to PD plate without antibiotic to get rid of CRISPR plasmid. The resulting conversion mutants have no antibiotic resistance. Correct integration of the recombinant DNA was confirmed by southern blot (not shown).

The conversion mutants and corresponding effector deletion mutants (Schilling *et al*., 2014) were used for infection. Disease scoring was done as previous described at 12 dpi. Disease indexes 9, 7, 5, 3, 1 and 0 were assigned to dead, heavy tumor, tumor, small tumor, chlorosis and normal symptom, respectively. The number of diseased plants were multiplied by the corresponding disease index, and the sum was divided by the total number of plants used for infection to give an average disease index. Student *t*-test was used for significance test of disease index from three biological replications.

## Results

### Maize leaf infection of *S. reilianum* and *U. maydis*

For a comparative transcriptomics approach between *S. reilianum* and *U. maydis*, we first tracked infection and fungal growth inside the maize leaves to record the milestone events during infection and define appropriate timepoints for analysis (Fig. **1**). 18-24 hours post infection, the compatible sporidia cells of both pathogens, mated on the leaf surface, and dikaryotic hyphae formed appressoria to penetrate the leaf surface (Fig. **1a**). At 2 days post infection (dpi), all infected leaves showed chlorosis as a first visible indication of successful infection (Fig. **S1a**). Microscopic analysis of WGA-AF488 stained hyphae showed similar colonization in the leaf vascular tissue for both smut fungi (Fig. **1b**). At 3 dpi, the earliest microscopic difference was observed between *S. reilianum* and *U. maydis*: in alignment to our previous report (Matei *et al*., 2018), host bundle sheath cells underwent *de novo* cell division in *U. maydis* infected leaves, indicating the initiation of tumorigenesis. In contrast, *S. reilianum* further accumulated in the leaf vein and showed aggregation as a string of round cells (Fig. **1c**). At 4 dpi, the first visible small tumor structures can be observed on *U. maydis* infected leaves, while *S. reilianum* infected leaves showed individual events of necrosis (Fig. S**1a**) and more frequently cell aggregation (Fig. **1d**, Fig. S**1b**). In parallel to the microscopic observation, fungal biomass in the infected leaves was quantified for both pathogens. The relative amount of both fungal pathogens increased similarly during infection until 3 dpi. Interestingly, at 4 dpi *S. reilianum* showed a significantly higher abundance compared to *U. maydis*, as the relative amount of *S. reilianum* doubled while the relative amount of *U. maydis* was maintained in comparison to 3 dpi (Fig. **1e**).

**Fig.1.**
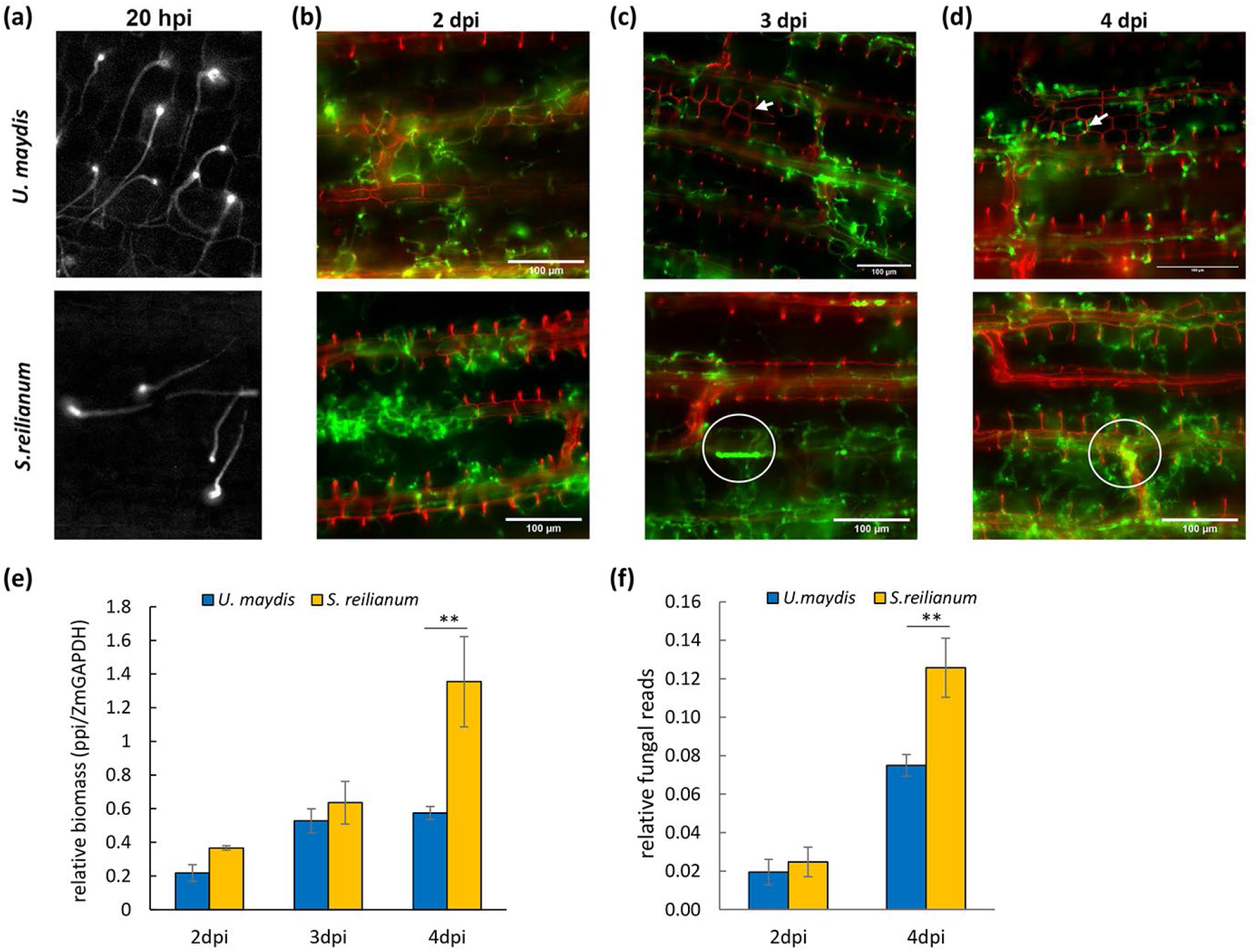
Comparison of *U. maydis* and *S. reilianum* growth during leaf infection. **a**, calcofluor white staining of fungal hyphae and appressorium at 20hpi. **b-d**, WGA-AF488-Propidium Iodide co-staining to show the fungal biotrophic growth inside the plant cell. Fungal cells were stained with WGA-AF488 (green), the vascular cell wall was stained with Propidium Iodide (red). White arrows indicate the *de-novo* divided bundle sheath cells triggered by *U. maydis* infection. White circles indicate the aggregation of *S. reilianum* cells inside the leaf vein. **e**, the relative biomass quantification by qPCR shows the fungal growth during infection, and a significant higher abundance of *S. reilianum* was detected at 4 dpi. **f**, the relative fungal reads from RNA-seq, which was the ratio between reads uniquely mapped to fungi and total reads uniquely mapped (fungi and maize). A significantly higher number of mapped reads was detected from *S. reilianum* at 4 dpi. “**”, Experiments have been performed in three independent biological replicates. *p*<0.05*. Student t-test* was used for significance test.

### Expression profiling of *U. maydis* and *S. reilianum* during maize leaf colonization

Based on our microscopic and biomass analysis, we decided to collect samples at 2 dpi to represent an infection stage where both *U. maydis* and *S. reilianum* successfully establish the biotrophic interaction with the host but do not show detectable differences in microscopical growth and biomass accumulation. Samples were also taken at 4 dpi when *U. maydis* tumorigenesis is initiated, while *S. reilianum* exhibits extensive tissue colonization without inducing morphological changes in the leaf. As control samples, cell pellets from axenic culture (AC) of both pathogens were used for RNA extraction. All samples have been generated in three independent biological replicates. In total, more than 670M paired-end reads were generated, of which over 93.7% of reads were uniquely mapped to either *U. maydis*, *S. reilianum* or the maize genome, respectively. The changes of fungal transcripts in the RNA-seq samples confirmed the biomass quantification results by qPCR. At 2 dpi, around 2% of fungal transcripts were mapped from both pathogens, and at 4 dpi, *S. reilianum* reads increased to 12.5% in the infected samples, which is significantly higher compared to *U. maydis* infected tissues (7.5%) (Fig. **1f**). Principal component analysis (PCA) analysis of *U. maydis* and *S. reilianum* samples showed that three biological replications from different conditions form distinct clusters, and variations between different conditions were larger than the variation of biological replications within the same condition (Fig. **2a**). To further analyze the RNA-seq data, we normalized reads counts and investigated the differentially expressed genes (DEGs) by edgeR (Robinson *et al*., 2009). After filtering the low- or not-expressed genes between samples, 6537 out of 6765 *U. maydis* genes and 6456 out of 6673 *S. reilianum* genes were used to identify DEGs. Compared to axenic culture, dramatic changes in gene expression were observed during seedling infection. Around 19.3-24.2% genes were up-regulated (log2 fold change >1, *p*<0.05) at 2 dpi and 4 dpi in both pathogens, while 12.1-19.1% genes were down-regulated (Fig. **2b**). We detected 435 out of 467 effector genes from *U. maydis* and 454 out of 489 from *S. reilianum* were expressed. As expected, most effector genes were specifically expressed in the biotrophic growth stage (311 in *U. maydis* and 307 in *S. reilianum*, respectively) and showed higher degrees of transcriptional induction compared to non-effector genes (Fig. **2c, d**), which reflects the explicit role of effectors in host infection. Three biological replicates of quantitative RT-PCR were conducted to confirmed the expression of several effector genes including infection markers such as Pit2 and Pep1 from our RNA-seq data (Fig. **S2a, b**).

**Fig.2.**
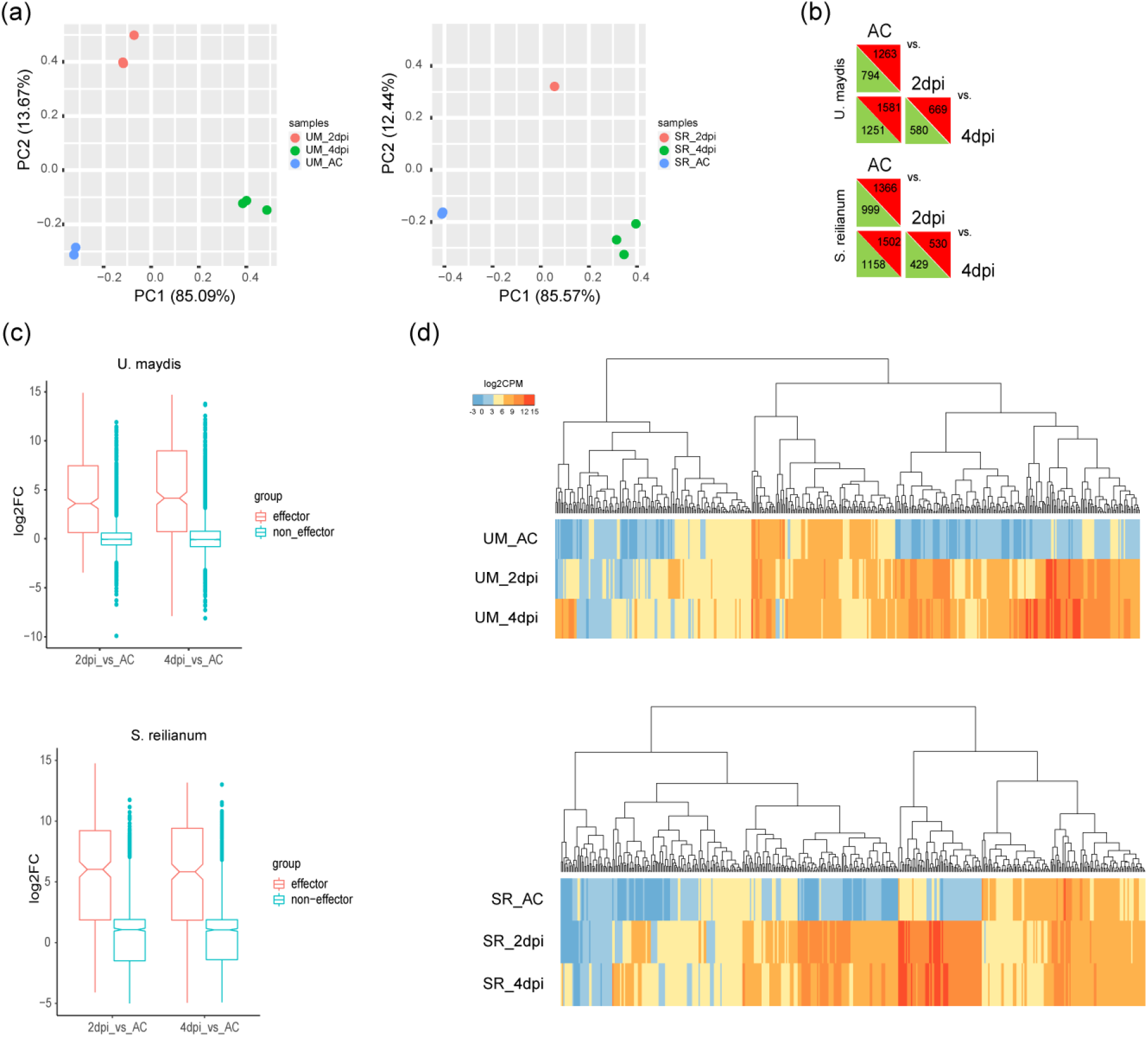
Overview of *U. maydis* and *S. reilianum* transcriptome, respectively. **a,** PCA analysis of *U. maydis* and *S. reilianum* RNA-seq samples. **b,** expression data from three different growth conditions. The number of differentially expressed genes (log2 fold change>1, p>0.05) from pairwise comparison were indicated in the triangles. Red and green triangles represent up-regulated and down-regulated genes between 2 dpi vs. axenic culture (AC), 4 dpi vs. AC and 4 dpi vs. 2 dpi, respectively. **c,** boxplot shows transcriptional changes of effector and non-effector genes during biotrophic growth. Effector genes were dramatically up-regulated compared to the non-effector genes. **d,** heatmap shows the overview of the expression changes of effector genes. “n” shows the number of effectors detected expressing in RNA-seq sample and the total number of effectors in the genome.

We clustered effector genes into all five possible expression patterns between three conditions (axenic culture, 2 dpi, 4 dpi) (Fig. **3a**). This classified groups of effector genes (i) showing constitutively increasing induction (pattern 1), (ii) induced *in-planta* on a stable level (pattern 2), (iii) being only transiently expressed (pattern3), (iv) dropping at 4 dpi (pattern 4), (v) or only being induced at 4 dpi (pattern 5). In *U. maydis* and *S. reilianum*, 227 effectors from each species were highly expressed across the whole biotrophic growth phases (pattern 1, 2 and 4), while only 14 *U. maydis* effectors and 8 *S. reilianum* effectors were transient induced at 2 dpi (pattern 3) and similar number of effectors were only required in the 4 dpi (23 from *U. maydis* vs. 25 from *S. reilianum* as pattern 5) (Fig. **3b**). The expression patterns are found to reflect previously observed virulence function of known effector genes. For example, the core effector *UmPep1*(*UMAG_01987*) (Doehlemann *et al*., 2009b; Hemetsberger *et al*., 2015) was stably expressed and clustered as pattern 2, while genes coding for the known virulence factors *UmPit2* (*UMAG_01375*) (Misas Villamil *et al*., 2019), *UmCmu1*(*UMAG_05731*) (Djamei *et al*., 2011), *UmTin2* (*UMAG_05302*) (Tanaka *et al*., 2014) and *UmRsp3* (*UMAG_03274*) (Ma *et al*., 2018) group to pattern 1 cluster. Interestingly, *S. reilianum* orthologs of these characterized effectors displayed different induction patterns, keeping a steady level between 2 and 4 dpi. In particular, we identified 100 out 467 of *U. maydis* effectors being induced in pattern 1, which is significantly higher compared to *S. reilianum* (43/488), whereas 184 out of 488 *S. reilianum* effector genes were showing stable expressing levels, which is higher as compared to *U. maydis* (136/467) (Fig. **3b**). This enrichment of different induction patterns of effector genes suggested different requirements in disease for the two species.

**Fig.3.**
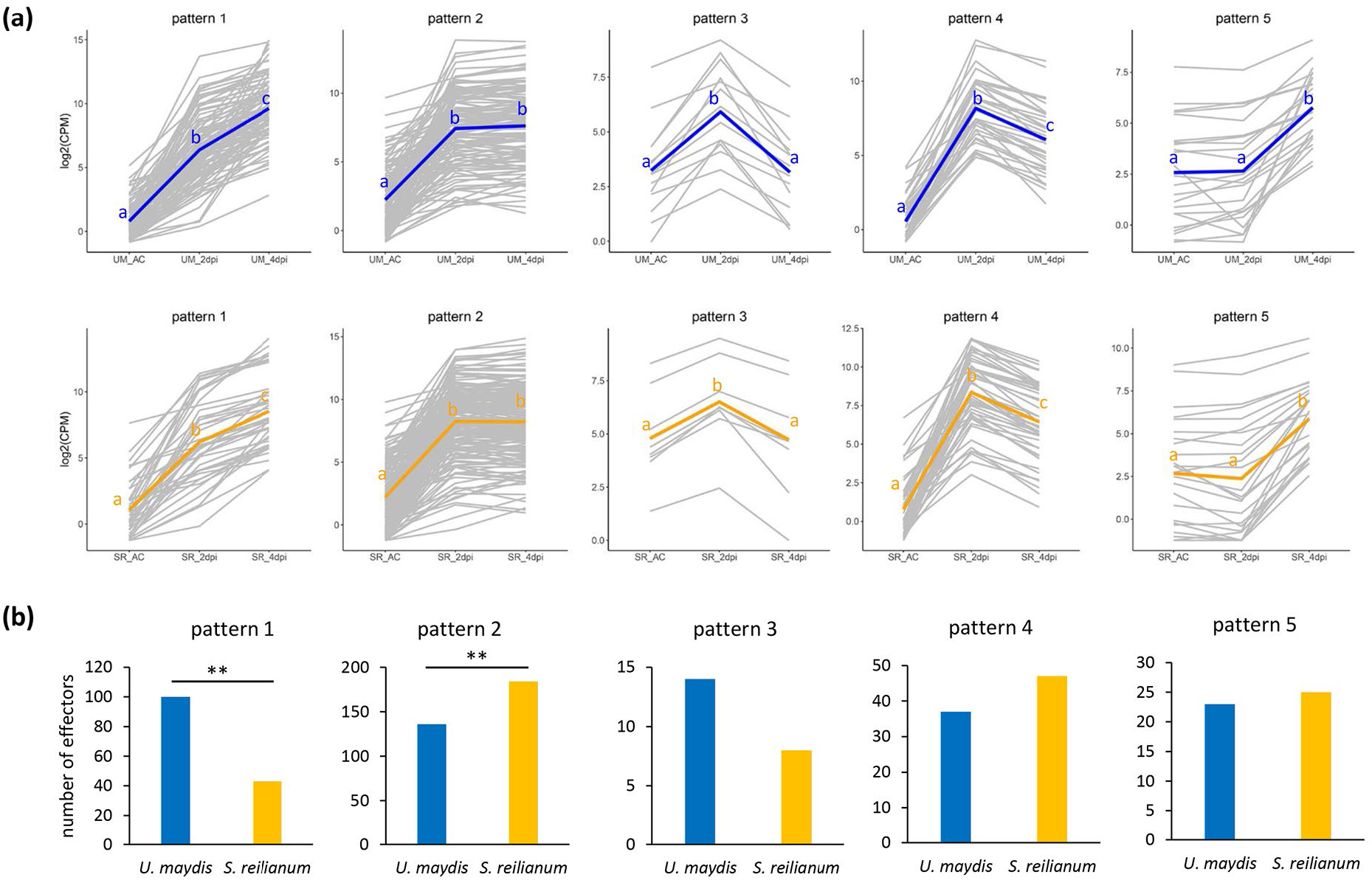
The induced pattern shows the expression changes of effectors. **a,** the induced effectors are clustered into 5 patterns based on the expression changes between axenic culture (AC), 2 dpi and 4 dpi conditions. The gray lines represent individual gene, and blue and orange lines are the mean of all genes in each cluster. “a”, “b” and “c” were used to indicate the significance levels between each condition. Genes with log2 fold change>1 and p<0.05 were considered significantly induced in pairwise comparison. **b,** the number of induced genes of each pattern. The *chi-square test* was used for statistic test for the significance. “**”, *p*<0.01.

### Differential regulation of effector orthologues in *U. maydis* and *S. reilianum*

To better understand how *U. maydis* and *S. reilianum* orchestrate their respective effector repertoires with respect to their differential disease development, we directly compared the relative expression levels of effector orthologs. We used orthoMCL to identify one-to-one ortholog pairs between *U. maydis* and *S. reilianum* and obtained 6005 one-to-one ortholog pairs including 335 effectors. This means that for avoiding ambiguous results due to functional redundancy or paralog compensation, one-to-many or many-to-many ortholog pairs were not included in this analysis. Relative expression was corrected for putative differences in length between the orthologs. PCA analysis based on the relative expression of 6005 ortholog pairs confirmed that plant associated samples form distinct clusters on time and species levels. In contrast, the axenic cultured samples of the different species have minor differences and clustered together (Fig. **4a**). Pearson correlation analysis displayed similar results and suggested the difference between samples increased at 4 dpi compared to 2 dpi, as the correlation decreased (Fig. S**3a**).

**Fig.4.**
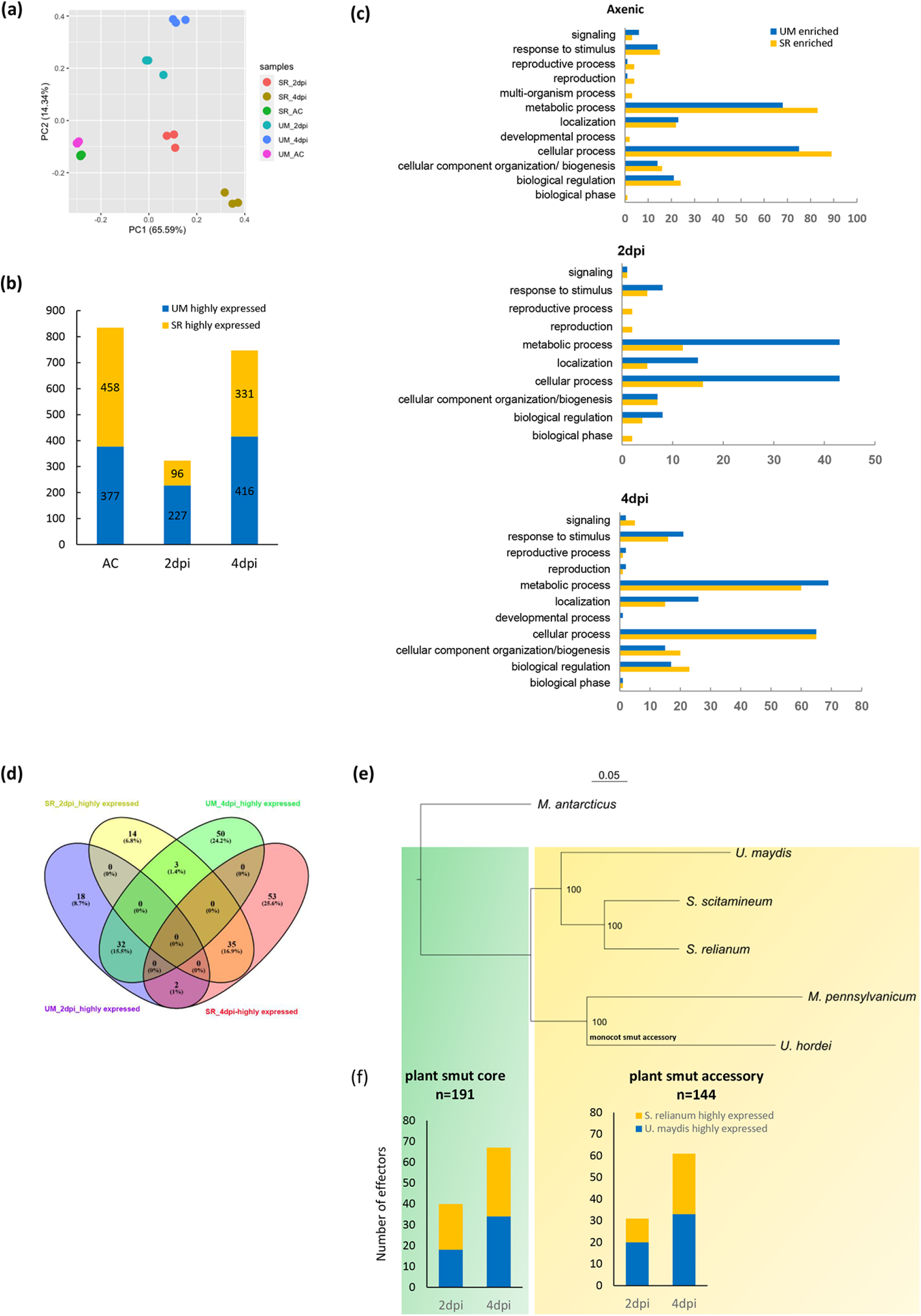
Orthologs were differentially regulated between *U. maydis* and *S. reilianum*. **a,** PCA analysis based on the relative ortholog expression of 6005 one-to-one ortholog pairs from *U. maydis* and *S. reilianum*. **b,** the number of DEOs detected in different conditions. The fold change>3 and p<0.05 were used to identify DEO. **c,** the GO classification of DEOs in three different conditions. **d,** the Venn diagram shows the number of differentially expressed effector orthologs during biotrophic growth. **e-f,** the differentially regulation of effectors clustered by the conservation between 5 smut fungi. **e,** the phylogenetic tree of 5 smut fungi, including *U. maydis*, *S. reilianum*, *Ustilago hordei*, *S. scitamineum* and *M. pennsylvanicum*. **f,** the number of effector DEOs in each category. *Chi-square test* was used for significance test. “**”, *p*<0.01, “***”, *p*<0.001.

Next, we compared the relative ortholog expression levels to identify <u>d</u>ifferentially <u>e</u>xpressed <u>o</u>rthologs (DEOs) in the two smut fungi. In total, we found 769 DEOs with comparatively higher expression in *U. maydis* and 737 DEOs being more highly expressed in *S. reilianum*, respectively (Fig. **4b**). The largest number of DEOs (835) was detected from axenic cultured samples (Fig. **4b**), and around 30 and 50% of these DEOs from *U. maydis* and *S. reilianum* were only confined in this stage (Fig. S**3b)**. The DEO number dropped to 323 at 2 dpi when the pathogens had switched from *in-vitro* yeast growth to the initial filamentous, biotrophic growth in the host tissue. At 4 dpi the number increased to 747, which likely reflects the significant differences in disease development between the two pathogens, i.e. initiation of tumorigenesis by *U. maydis* versus extended proliferation of *S. reilianum* for systemic spreading. We also detected more *U. maydis*-induced DEOs during infection, particularly at 2 dpi, when 227 DEOs were *U. maydis*-induced but only 97 DEOs for *S. reilianum* (Fig. **4b**). However, we could not detect overrepresented gene ontology terms, except for 4 dpi, where an enrichment of cellular lipid catabolic process (GO:0044242) was detected in the higher expressed *U. maydis* DEOs. The DEOs were involved in different biological processes related to pathogen growth (Fig. **4c**), suggesting that during the yeast and biotrophic growth no particular cellular or metabolic pathway was selected to promote cell growth in either *U. maydis* or *S. reilianum*.

From the 335 one-to-one effector orthologs, more than 60% were differentially regulated, including 100 *U. maydis*- and 102 *S. reilianum*-highly expressed effectors, from which 32 and 35 showed consistently enhanced expression during the two biotrophic time points (Fig. **4d**). The number of effector DEOs also increased from 99 at 2 dpi to 170 at 4 dpi as disease developed. Previously identified pathogenesis associated TFs might be responsible in the regulation of effector DEOs, particularly at 4 dpi, since differential expression was found for virulence related TFs *Rbf1*, *Biz1*, *Hdp2*, *Fox1* and *Nlt1* (Fig. **S4**). TF *Rbf1* was highly induced in *S. reilianum*, especially at 4 dpi, the transcript levels were 17-fold higher than those in *U. maydis*, which may lead to the high abundance of *Biz1* in *S. reilianum* (Fig. **S4**). On the other hand, the expression level of TFs *Hdp2*, *Fox1* and *Nlt1* were significantly higher in *U. maydis* at 4 dpi, which could indicate that these TFs play a role in the regulation of effectors highly expressed in *U. maydis* during tumorigenesis (Fig. **S4**). Besides these, the TF *ROS1* showed similar expression level in both species, which may reflect its dedicated function in sporogenesis, a process which needs to be initiated in both pathogens for completing the pathogenic like cycle (Fig. **S4**). The expression of *ROS1* is dramatically increased since 6 dpi in *U. maydis* that is corresponding to the initiation of sporogenesis (Tollot *et al*., 2016), which may explain why we did not observe its differential expression between two species. Cluster 19A (Fig. **S5a**) is the largest effector cluster, which is a crucial determinant of virulence in both two pathogens (Brefort et al., 2014; Ghareeb et al., 2019). We observed that 10 out of 14 effectors residing in cluster 19A were *U. maydis*-enhanced, whereas only two were highly expressed in *S. reilianum*, including the neofunctionalized Tin2 (Tanaka *et al*., 2019) (Fig. **S5b**).

In a previous study, Schuster et al. analyzed 12 smut related basidiomycete and suggested that effector repertoires of smut fungi comprise sets of core and accessory effectors (Schuster *et al*., 2018). Based on this information, we classified the 335 one-to one effector pairs into plant smut core effectors, which were present in all 4 monocot smut fungi, including *Ustilago hordei* and *Sporisorium scitamineum*, and accessory effectors, which present in *U. maydis* and *S. reilianum* but not in all other plant smut fungi (Fig. **4e**). *Melanopsychium pennsylvanicum* was included in this analysis due to its close phylogenetic relationship to monocot smut pathogens, which suggests a recent host jump event (Sharma et al. 2014). In total, we identified 191 core effectors and 144 accessory effectors. At 2 dpi, 40 core effectors and 31 accessory effectors were differentially expressed between *U. maydis* and *S. reilianum*, and these numbers were significantly increased to 67 and 61 at 4 dpi, respectively (Fig. **4f**). We did not find accessory effectors subjected to more intensive transcription regulation compared to core effectors (Fig. **4f**), suggesting that differential regulation of both core and accessory effectors is needed for the distinct pathogenic development in the two pathogens.

### Functional conservation and neofunctionalization of differentially expressed effector orthologs

Several *U. maydis* effectors have been functionally characterized, which provides a reference to link the ortholog expression with their biological functions. The conserved core effector Pep1 (Hemetsberger et al., 2012, 2015; Sharma et al. 2019), as well as the virulence factors Pit2 (Misas Villamil *et al*., 2019) and Cmu1 (Djamei *et al*., 2011) were highly induced during biotrophic growth (Table. **S2**) and showed similar expression level between *U. maydis and S. reilianum*, which is consistent with their crucial roles in maize immunity inhibition (Fig. **5a**). The effectors Rsp3 (Ma *et al*., 2018) and ApB73 (Stirnberg & Djamei, 2016) had been validated in previous studies for their conserved function between two smut fungi by ortholog complement assay, and also these genes show comparable expression levels (Fig. **5a**). Also the See1 effector, which in *U. maydis* has a virulence function specifically for leaf tumor-formation (Redkar *et al*., 2015a), is expressed in *S. reilianum* both at 2 and 4 dpi (Fig. **5a**). Another effector DEO which caught our attention is Tin2. SrTin2 neofunctionlized during speciation between *U. maydis* and *S. reilianum* (Tanaka *et al*., 2019). We found a significantly enhanced expression of *SrTin2* compared to *UmTin2* (Fig. **5a**), in line with the neofunctionalisation, as the transcriptional level divergence could be involved to promote the functional divergence.

**Fig.5.**
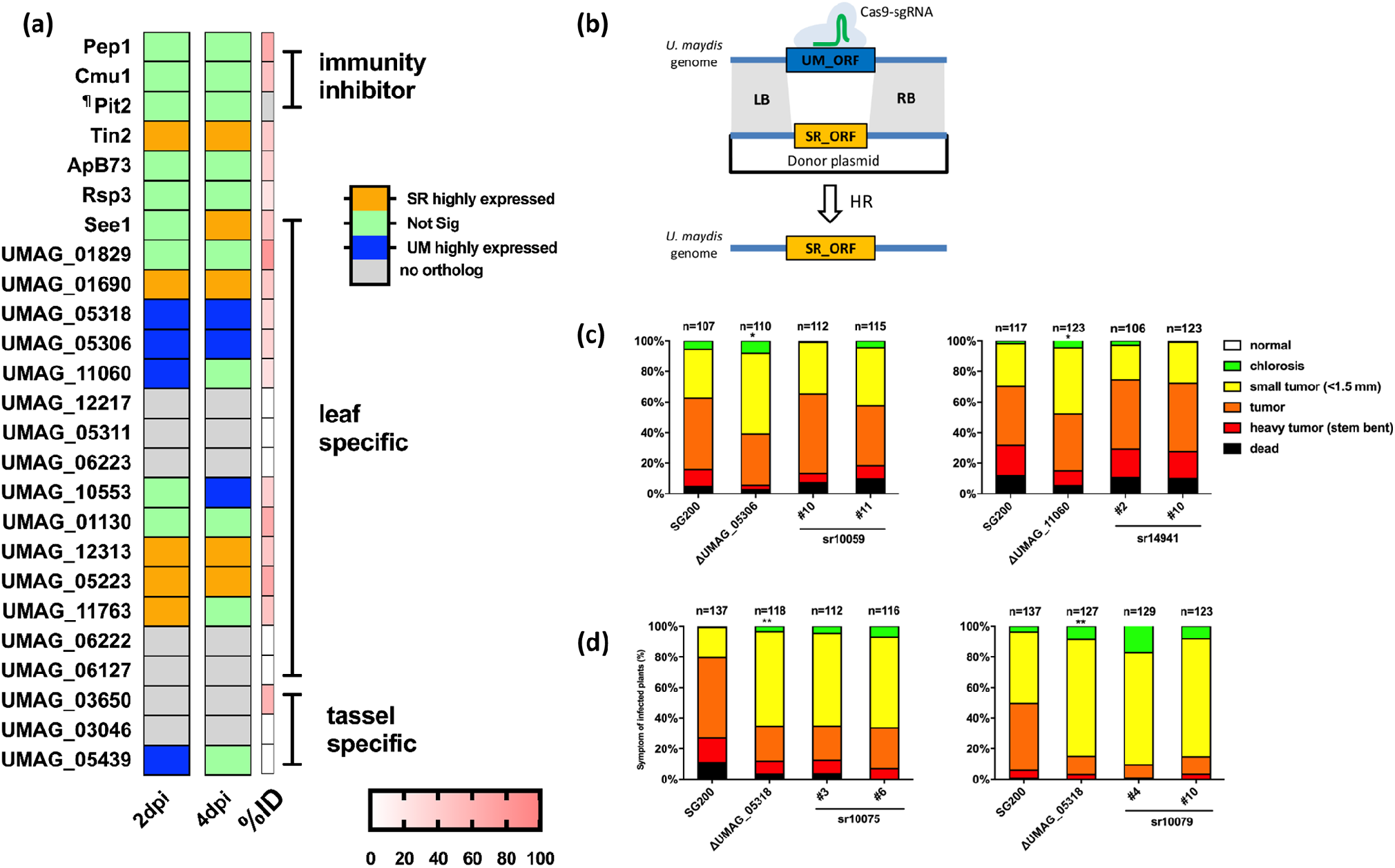
Effector orthologs between *U. maydis* and *S. reilianum* are functional conserved. **a,** a heatmap shows the expression difference and protein sequence identity of effectors ortholog pairs which have been knocked out and subjected to virulence tests in *U. maydis*. “¶”, Pit2 only conserved in functional PID motif which failed to be detected during ortholog identification and “%ID” was not showed. **b,** the scheme demonstrated the process of CRISPR-Cas9 mediated gene conversion. The sgRNA was designed to target the coding sequence of *U. maydis* effector, and the donor plasmid contains *S. reilianum* ortholog coding sequence flanked by 1kb homologous region. **c-d,** Disease scoring results of gene conversion mutants and corresponding *U. maydis* knock out mutant. **c,** Converted UMAG_05306 and UMAG_11060 to the *S. reilianum* orthologs did not affect the virulence in SG200. **d,** Converted UMAG_05318 to its *S. reilianum* ortholog Sr10075 or Sr10079, the paralog of Sr10075, compromise SG200 virulence, and show reduced virulence as deletion mutant. Experiments have been performed in three independent biological replicates. n indicates the total number of plants tested in three replicates. The significance as calculated by *student t-test* with disease index “*”, *p*<0.05, “**”, *p*<0.01.

From a set of 20 organ-specific effectors, we found 9 effector DEOs and 8 of them were leaf-specific effectors, from which 4 were validated as virulence factors from a deletion mutant screen (Schilling *et al*., 2014) (Fig. **5a**). Orthogroups *UMAG_11060/Sr14941*, *UMAG_05306/Sr10059* and *UMAG_05318/Sr10075* were particularly interesting as they were induced in both pathogens (Table. **S2**), however, the transcripts levels of the orthologs in *U. maydis* were at least 4.8 folds (for orthogroup *UMAG_11060/Sr14941* at 2 dpi) to more than 50 times (orthogroup *UMAG_05318/Sr10075* at 4 dpi) higher compared to their *S. reilianum* orthologs (Table. S2). These effectors are specifically required for *U. maydis* induced leaf tumors (Schilling *et al*., 2014). Thus, their low expression in *S. reilianum* might suggest that these genes are of lesser importance for the basal establishment of biotrophic growth. We assessed the protein function of these DEOs to test if functional diversification occurred and is associated with differentially expression, as in the case of Tin2. We converted the open reading frame of the candidate effectors in *U. maydis* to their respective *S. reilianum* orthologs. For the generation of *in situ* seamless mutants we used a CRISPR-Cas9 approach that allowed marker-free selection of gene-replacement mutants in *U. maydis* (Fig. **5b**). In each case, an sgRNA was designed to target the effector gene to induce a double strand break, which was then repaired by HDR with a donor plasmid containing the *S. reilianum* effector ortholog, flanked by 1kb homologs arms (Fig. **5b**). This CRISPR-Cas9 mediated gene conversion eliminated potential ectopic effects and allowed expression of the orthologs under the native *in-situ* cis- or trans-regulatory elements.

In maize infection assays, conversion of UMAG_05306 and UMAG_11060 to their *S. reilianum* orthologs did not affect the virulence compared to the *U. maydis* progenitor strain SG200, while the deletion of these two effectors resulted in significantly reduced virulence (Fig. **5d**). However, conversion of UMAG_05318 into its *S. reilianum* ortholog Sr10075 significantly reduced virulence to the similar level of the deletion mutant, which indicates that Sr10075 might have a different function, or lost its activity during speciation (Fig. **5d**). In *S. reilianum*, Sr10075 has a paralog Sr10079, which has only low sequence identity to UMAG_05318 and therefore was not detected in our one-to-one ortholog analysis (Fig. **S6a, b**). Deletion of Sr10079 was previously shown to reduce the virulence in *S. reilianum* (Ghareeb *et al*., 2019). Conversion of UMAG_05318 into Sr10079 resembled virulence defect of the UMAG_05318 deletion mutant. Thus, neither Sr10075, nor Sr10079 can functionally replace UMAG_05138, suggesting the functional divergence of the orthogroup *UMAG_05318/Sr10075 - Sr10079* during speciation, which identifies them as an interesting objective to test for possible neofunctionalization.

To further confirm and investigate how promoters regulate the expression of DEOs, we used the CRIPSR-Cas9 mediated conversion approach to replace the promoter of the *U. maydis* induced DEOs *UMAG_11060* and *UMAG_05306* with their respective *S. reilianum* counterparts in SG200 (Fig. **S7a**), and quantitative RT-PCR was used to detect the chimeric gene expression. Infected samples at the time points when the differential expression levels were detected in RNA-seq (2 dpi for *UMAG_11060* and 4 dpi for *UMAG_05306*, Fig. **5a**) were tested. Expressing *UMAG_11060* under Pro^sr14941^ showed lower expression at 2 dpi (Fig. **S7b**), which was similar to our RNA-seq analysis, indicating the cis-regulation in the promoter region determined their differential expression. However, Pro^Sr10059^ surprisingly drove a similar expression level of *UMAG_05306* at 4 dpi in *U. maydis* genetic background, suggesting a trans regulator or distal enhancer, which is variable between two species control the differential expression (Fig. **S7b**).

Together, the results of the gene conversion approach showed that functional diversification of effector orthologs can be caused on two levels: (i) the differential expression of orthologous genes which encode functionally conserved proteins (as found for orthogroups UMAG_05306/Sr10059 and UMAG_11060/Sr14941), (ii) the functional divergency of effectors on the protein level, which further fine-tuned on transcription levels for functional adaption (iii) the regulations of DEO is complicated, involving both cis- and trans-regulatory elements.

## Discussion

In this study we applied a cross-species transcriptome comparison between *U. maydis* and *S. reilianum* to investigate transcriptomic changes of effector orthologs during maize leaf infection.

A crucial challenge in cross-species RNA-seq is to determine the right timing for sampling to exclude that the observed differences result rather from the different disease progression states than from a differential regulation as such. Sucher *et al*. (2020) collected samples from the edge of disease lesions, which were smaller than 25 mm in diameter, as indication of comparable infection stage to study the different host responses to *Sclerotinia sclerotiorum* for quantitative disease resistance (Sucher *et al*., 2020). Time resolved microscopes were applied in *Fusarium virguliforme* to facilitate RNA-seq to study its transcription plasticity on different hosts (Baetsen-Young *et al*., 2020). In our experiment, we conducted a series of microscopy combined with qPCR-based biomass quantification, and selected two biotrophic interaction phases representing a comparable (2 dpi) and a distinct colonization stage (4 dpi) for RNA-seq.

When comparing the induction profiles of effectors in *U. maydis* and *S. reilianum* respectively, we found a significantly higher number of *S. reilianum* effectors showing stable expression between 2 and 4 dpi, while in *U. maydis* more effectors displayed an increasing expression over time. We further focused on the expression regulations of one-to-one effector orthogroups. Overall, 207 out of 335 one-to-one effector orthologs were differentially expressed between *U. maydis* and *S. reilianum* during biotrophic interaction, and the number of effector DEOs increased at 4 dpi. The difference in effector induction patterns and differential regulation of effector DEOs are consistent with the appearance of different symptoms. At 4 dpi, *U. maydis* triggers local host cell proliferation to produce tumors, which will require the high transcript abundance of effector genes related to tumorigenesis; in contrast, *S. reilianum* sustaining stable expression levels of effectors likely reflects maintenance of the biotrophic interaction in already colonized tissue and continuation of systemic proliferation through the leaf veins with inducing structural changes in the host tissue

Not only effector genes, but also pathogenesis related transcription factors were found to be differentially regulated, which in turn is probably playing a crucial role in the control of effector DEO expression. TFs Rbf1 and Biz1 were highly expressed in *S. reilianum*, which may be responsible for the expression of *S. reilianum* enriched effector DEOs. However, the abundance of TF Hdp2 transcripts, which is activated by Rbf1 (Heimel *et al*., 2010b), was more highly expressed in *U. maydis*, indicating the complicated regulation of TF cascade. The promoter conversion of Pro^Sr10059^ showed comparable expression with Pro^UMAG_05306^ in *U. maydis* (Supplementary Figure S7), indicating the potential trans effects such as transcription factors or distal enhancers might be driving differential regulation while for orthogroup UMAG_11060/sr14941, cis-regulatory elements in promoter region likely determines the expression pattern. Until now, only few direct target genes of TFs were identified by CHIP-seq in smut fungi (Tollot *et al*., 2016). Elucidation of the binding motif of these pathogenesis related TFs and combined with the promoter analysis of effector DEOs, which were obtained in this study could be a starting point for a more detailed study on the specific roles of TFs in the orchestration of effector expression with respect to the adaptation to different infection styles in pathogen speciation.

A major conclusion which can be drawn from the comparative transcriptome analysis is that the expression levels of effector orthologs between *U. maydis* and *S. reilianum* are related to their function: effectors with general virulence functions showed similar expression levels between both smut fungi, whereas effectors adapted to execute specific processes such as the leaf-tumor specific effectors of *U. maydis* showed significant differences in their transcriptional regulation. UMAG_05306 and UMAG_11060, whose disruption resulted in reduced virulence in *U. maydis* and were functionally conserved with respect to the corresponding *S. reilianum* orthologs. This implies that the functional diversification of effectors can be driven on the transcriptional level and is not necessarily linked to a changed protein function. However, UMAG_05318 provides another example for combination of transcriptional plasticity and functional diversification of the protein level. While the orthogroups UMAG_05318/Sr10075 or Sr10079 and Umtin2/Srtin2 were expressed during leaf infection both in *U. maydis* and *S. reilianum*, they also show similar induction pattern during biotrophic interaction. However, when comparing the expression levels of the orthologs, UMAG_05318 is more than 50 times higher than its *S. reilianum* ortholog, while Srtin2 has a 10 times higher expression compared to its respective ortholog in *U. maydis*. Until now, the molecular mechanisms of these differentially regulated organ specific effectors are not elucidated in *U. maydis*, and whether they also contribute to virulence in *S. reilianum* is largely unknown. Deletion of Sr10075 alone did not seem affect virulence, while a knock-out Sr10079 reduced *S. reilianum* virulence (Ghareeb *et al*., 2019). The orthologs of UMAG_05318 were found in *S. scitamineum* and *S. reilianum f. sp. reilianum*, but not in *U. hordei* and *M. pennsylvanicum*. Elucidation of the function of this ortholog group and its role in smut fungi evolution and speciation will be our next aim. Interestingly, both effectors showing functional divergence are located in effector cluster 19A, which is the largest virulence cluster in both pathogens (Brefort *et al*., 2014; Ghareeb *et al*., 2019) and several *U. maydis* highly expressed effector DEOs were also identified in this cluster from our study. However, the impact of 19A deletion on *S. reilianum* virulence was monitored by the head smut phenotype in the maize inflorescence (Ghareeb *et al*., 2019). Thus, it could be possible that these effectors were highly expressed in a different time/spatial pattern to contribute to the systemic spreading of *S. reilianum*. On the other hand, it would be tempting to study if higher expression of cluster 19A effectors could promote the adaptation of *S. reilianum* to leaf infection and eventually even trigger the formation of leaf tumor by this pathogen. Therefore, to elucidate the molecular functions of these DEOs and the identification of their host targets will be instrumental to gain new insights of how evolution of effector orthologs promote the emergence of new pathogens.

With only a limited set of virulence DEOs can be tested in gene conversion experiments, this study identified three effector DEOs as promising candidates for effectors with virulence function in speciation. Despite these findings, one should not neglect the importance of effectors that evolved new functions without fundamental changes in expression patterns. For this, the Pit2 effector is a potential example: SrPit2 showed a similar expression level compared to UmPit2. However, in a previous study we found that SrPit2 only partially complemented a *U. maydis* pit2 knock out mutant, which was linked with a reduced ability of SrPit2 to inhibit SA-associated PLCPs (Misas Villamil *et al*., 2019). Furthermore, a recent apoplastic proteome analysis suggested that composition of PLCPs is different between maize leaf and root (Schulze Hüynck *et al*., 2019). Given that in natural conditions, *S. reilianum* infects maize through the root while *U. maydis* infects aerial tissues, one could speculate that Srpit2 adapted to be a more efficient inhibit of root PLCPs, while UmPit2 adapted to inhibit PLCPs in the maize leaf. This reflects that effector function can specialize to different hosts of tissues. In addition to this mechanism of gradual effector specialization, our approach to compare the transcriptional levels of orthologs across two different pathogen species documented that both the transcriptional plasticity and functional divergency of effector proteins contribute to pathogen speciation. Including more phylogenetically related smut fungi infecting different hosts, such as *S.□reilianum f. sp. reilianum* for sorghum *and U. hordei* for barley in future research will provide a more comprehensive map of the emergence and adaptation of pathogenicity effectors during host adaptation and specialization. Besides the plastic regulation of effector orthogroups, one should not ignore the role of species-specific effectors in virulence, which is not discussed in our study. Studies on how pathogens organize their effector warehouse in temporal and spatial manner to shape the pathogenesis, and how these effectors were adapted during speciation will shed a light on the evolution and mechanistic basis of plant-pathogen interactions.

## Acknowledgments

This work was funded by the European Research Council under the European Union’s Horizon 2020 research and innovation program (consolidator grant conVIRgens, ID 771035), as well as funding by the Deutsche Forschungsgemeinschaft (DFG, German Research Foundation) under Germany’s Excellence Strategy-EXC-2048/1-Project ID: 390686111. Weiliang Zuo and Jasper Depotter are supported by the Research Fellowship Programme for Postdoctoral Researchers of the Alexander von Humboldt Foundation. MT and DKG received support by the LOEWE initiative of the government of Hesse, in the framework of the LOEWE Centre for Translational Biodiversity Genomics (TBG) and the LOEWE Cluster for Integrative Fungal Research (IPF).

## Author contributions

W.Z. and G.D. designed the research; W.Z. performed the data analysis, molecular experiment and virulence assay; J.D. and D.K.G. mapped the RNA-seq data; J.D. conducted phylogenetic analysis; D.K.G. and M.T. provided ortholog expression normalization; W.Z. and G.D. wrote the paper with contributions from the other authors.

## Data availability

RNA sequencing data has been submitted to NCBI Genbank and are available under the following accession: BioProject ID PRJNA674747

## Supplementary Captions

**Supplementary Fig.1 The symptom of *U. maydis* and *S. reilianum* infected leaves. a**, The photos shows the symptom development from 2 dpi to 4 dpi. The dashed circle shows the first visible tumor from *U. maydis* infected leaf while the dashed rectangle shows the necrosis spot caused by *S. reilianum* infection. The back *of U. maydis* infected leaf from 4 dpi is shown to better visualize the tumor. **b,** Confocal microscopy shows the details of the *S. reilianum* aggregation inside the leaf vascular at 4 dpi after WGA-AF488-Propidium Iodide staining. The fungal cells aggregate inside the vascular tissue. Scale bar=100 μm.

**Supplementary Fig.2 qPCR confirms the expression of effector genes from *U. maydis* and *S. reilianum*. a,** Expression levels of *U. maydis* effector genes. **b,** Expression levels of *S. reilianum* effector genes. Infection markers such as *UmPep1* and *UmPit2* and their corresponding orthologs were detected, as well as organ-specific effector See1 and two differentially expressed orthogroups *UMAG_05306/Sr10059* and *UMAG_11060/Sr14941*. The relative expression levels of effectors were normalized by respective reference *peptidylprolyl isomerase* (*ppi*) genes by 2^-ΔCt^ method. Error bar indicate standard deviation from three biological replications.

**Supplementary Fig.3 Differential regulation of orthologs between *U. maydis* and *S. reilianum* from different samples. a,** Pearson correlation analysis between *U. maydis* and *S. reilianum* samples based on relative ortholog expression from 6005 one-to-one orthologs. The *r* values indicate a high to low correlation between *U. maydis* and *S. reilianum* samples: AC>2 dpi>4 dpi. **b,** Venn diagram shows the number of differentially expressed ortholog in *U. maydis* and *S. reilianum* from different conditions. About 30% and 50% of detected DEOs were only detected from axenic culture.

**Supplementary Fig.4 Expression of pathogenic development related transcription factors between *U. maydis* and *S. reilianum* during biotrophic growth from RNA-seq data.** The transcription factors *Rbf1* and *Biz1* were highly expressed in *S. reilianum* while transcription factors *Hdp2*, *Fox1* and *Nlt1* were enhanced expressed in *U. maydis*. The transcription factor *Ros1* shows similar expression between these two smut fungi. *Student t-test* was used for significance test. “**”, *p* <0.01.

**Supplementary Fig.5 The distribution of effector DEOs on chromosomes. a,** the distribution of DEOs in 22 chromosomes based on the *U. maydis* genome organization. The blue bar shows *U. maydis* highly expressed DEOs, and the yellow bar shows the *S. reilianum* highly expressed DEOs. 2 dpi DEOs and 4 dpi DEOs were plotted above and below the chromosome, respectively. The red rectangle shows effector cluster 19A. **b,** the diagram shows the overview of synteny of effectors in cluster 19A. The effector genes are showed in green arrows; non-effector genes are shown in dark gray. The synteny of effector genes are in grey shade, the outline color of the shade indicates the DEO. The *U. maydis* highly expressed effector ortholog pairs are outlined in blue, the *S. reilianum* highly expressed genes are outlined in yellow.

**Supplementary Fig.6 Sequence comparison of UMAG_05318, Sr10075 and Sr10079. a,** the maximum likelihood phylogenetic tree of UMAG_05318, Sr10075 and Sr10079 based on the amino acid sequences. Sr10079 is distant from UMAG_05318 and Sr10075. **b,** the amino acid sequence alignment of UMAG_05318, Sr10075 and Sr10079. The identical amino acids were shaded in grey.

**Supplementary Fig. 7 The regulation of *S. reilianum* promoters in *U. maydis* by CRIPSR-Cas9 mediated promoter conversion. a,** Schematic model of the method used for CRISPR-Cas9 mediated promoter conversion. **b,** qPCR detection of the corresponding effectors expression under different orthologous *S. reilianum* promoters. At 4 dpi, Pro^sr10059^ regulates gene expression to the similar level as Pro^UMAG_05306^ in *U. maydis*, while at 2 dpi, Pro^sr14941^ drove gene expression was significantly lower compared to Pro^UMAG_11060^. Error bar indicates standard deviation from three biological replications.

**Supplement Table.1 Oligo sequences used.**

**Supplement Table.2 Log2 CPM (count per million) of fungal reads.**

**Supplement Table.3 Relative ortholog expression data of 6005 one-to-one orthologs between *U. maydis* and S. *reilianum***.

## Notes

### Competing Interest Statement

The authors have declared no competing interest.

