## Supplementary Figures for "Cross-species analysis between the maize smut fungi *Ustilago maydis* and *Sporisorium reilianum* highlights the role of transcriptional plasticity of effector orthologs for virulence and disease"

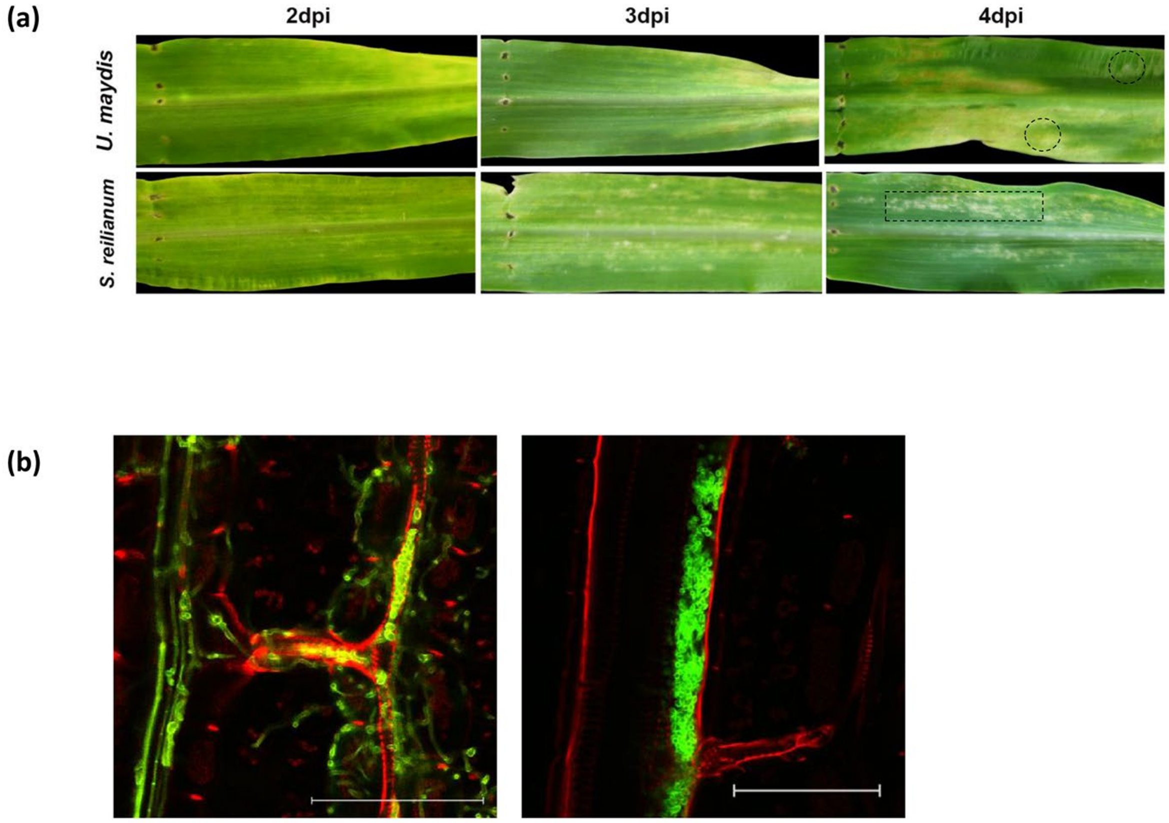


**Supplementary Fig.1 The symptom of *U. maydis* and *S. reilianum* infected leaves. a**, The photos shows the symptom development from 2 dpi to 4 dpi. The dashed circle shows the first visible tumor from *U. maydis* infected leaf while the dashed rectangle shows the necrosis spot caused by *S. reilianum* infection. The back *of U. maydis* infected leaf from 4 dpi is shown to better visualize the tumor. **b,** Confocal microscopy shows the details of the *S. reilianum* aggregation inside the leaf vascular at 4 dpi after WGA-AF488- Propidium Iodide staining. The fungal cells aggregate inside the vascular tissue.  Scale bar=100 µm.

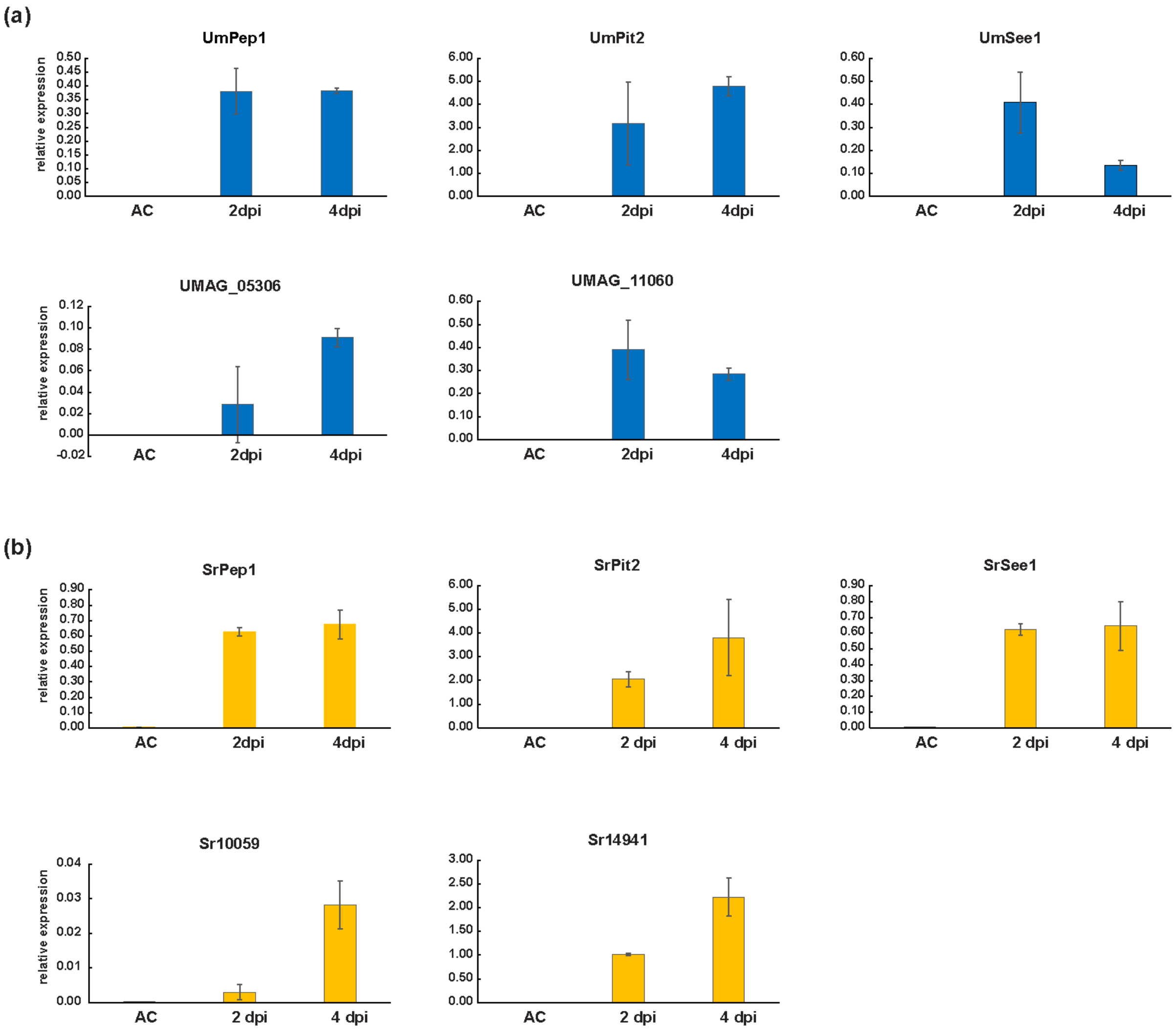


**Supplementary Fig.2 qPCR confirms the expression of effector genes from *U. maydis* and *S. reilianum*. a,** Expression levels of *U. maydis* effector genes. **b,** Expression levels of *S. reilianum* effector genes. Infection markers such as *UmPep1* and *UmPit2* and their corresponding orthologs were detected, as well as organ-specific effector See1 and two differentially expressed orthogroups *UMAG_05306/Sr10059* and *UMAG_11060/Sr14941*. The relative expression levels of effectors were normalized by respective reference *peptidylprolyl isomerase* (*ppi*) genes by $2^{-\Delta Ct}$ method. Error bar indicate standard deviation from three biological replications.

**(a)**

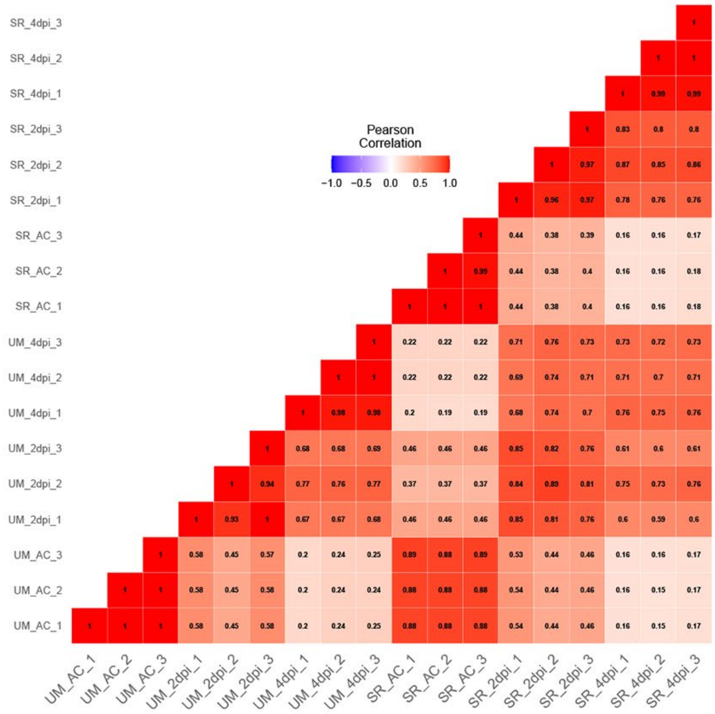


**(b)**

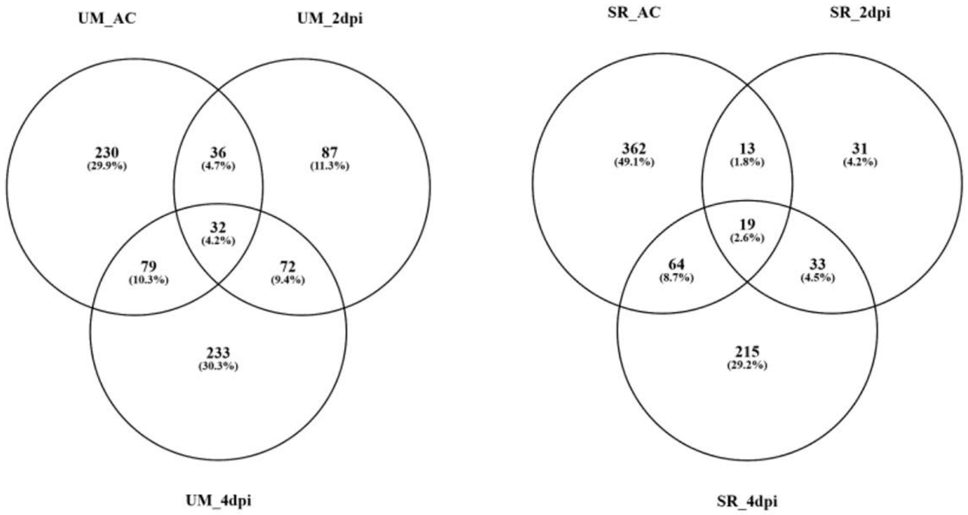


**Supplementary Fig.3 Differential regulation of orthologs between *U. maydis* and *S. reilianum* from different samples. a,** Pearson correlation analysis between *U. maydis* and *S. reilianum* samples based on relative ortholog expression from 6005 one-to-one orthologs. The *r* values indicate a high to low correlation between *U. maydis* and *S. reilianum* samples: AC>2 dpi>4 dpi. **b,** Venn diagram shows the number of differentially expressed ortholog in *U. maydis* and *S. reilianum* from different conditions. About 30% and 50% of detected DEOs were only detected from axenic culture.

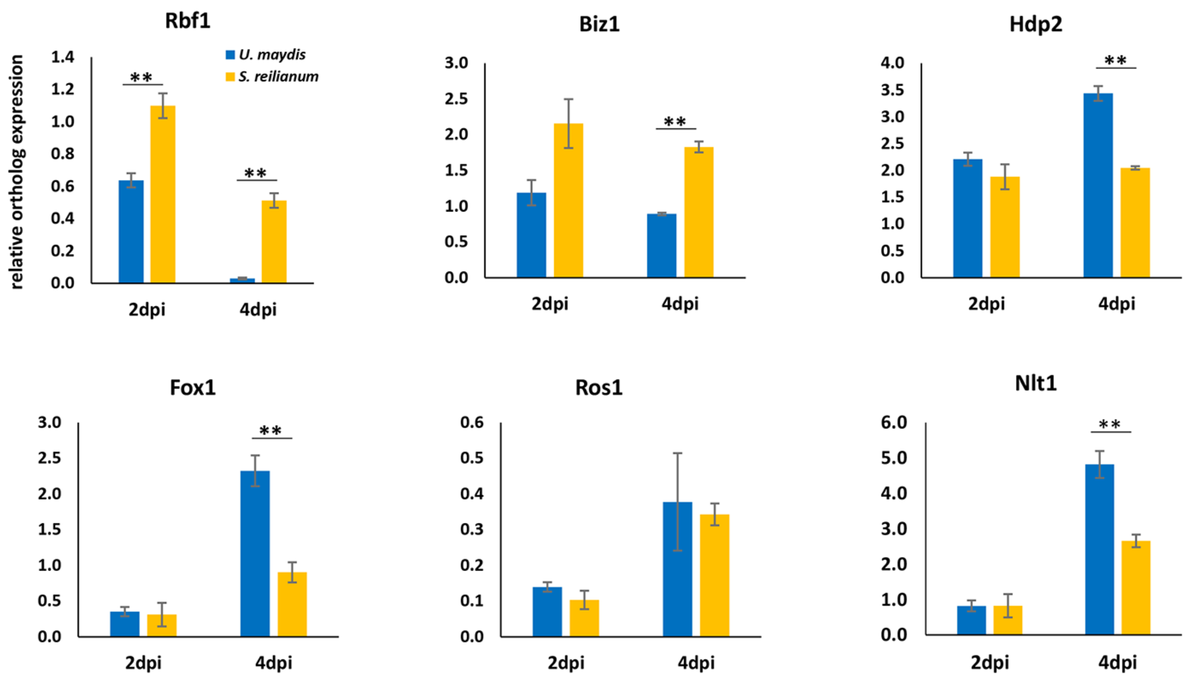


**Supplementary Fig.4 Expression of pathogenic development related transcription factors between *U. maydis* and *S. reilianum* during biotrophic growth from RNA-seq data.** The transcription factors *Rbf1* and *Biz1* were highly expressed in *S. reilianum* while transcription factors *Hdp2*, *Fox1* and *Nlt1* were enhanced expressed in *U. maydis*. The transcription factor *Ros1* shows similar expression between these two smut fungi. *Student t-test* was used for significance test. "**", $p$ <0.01.

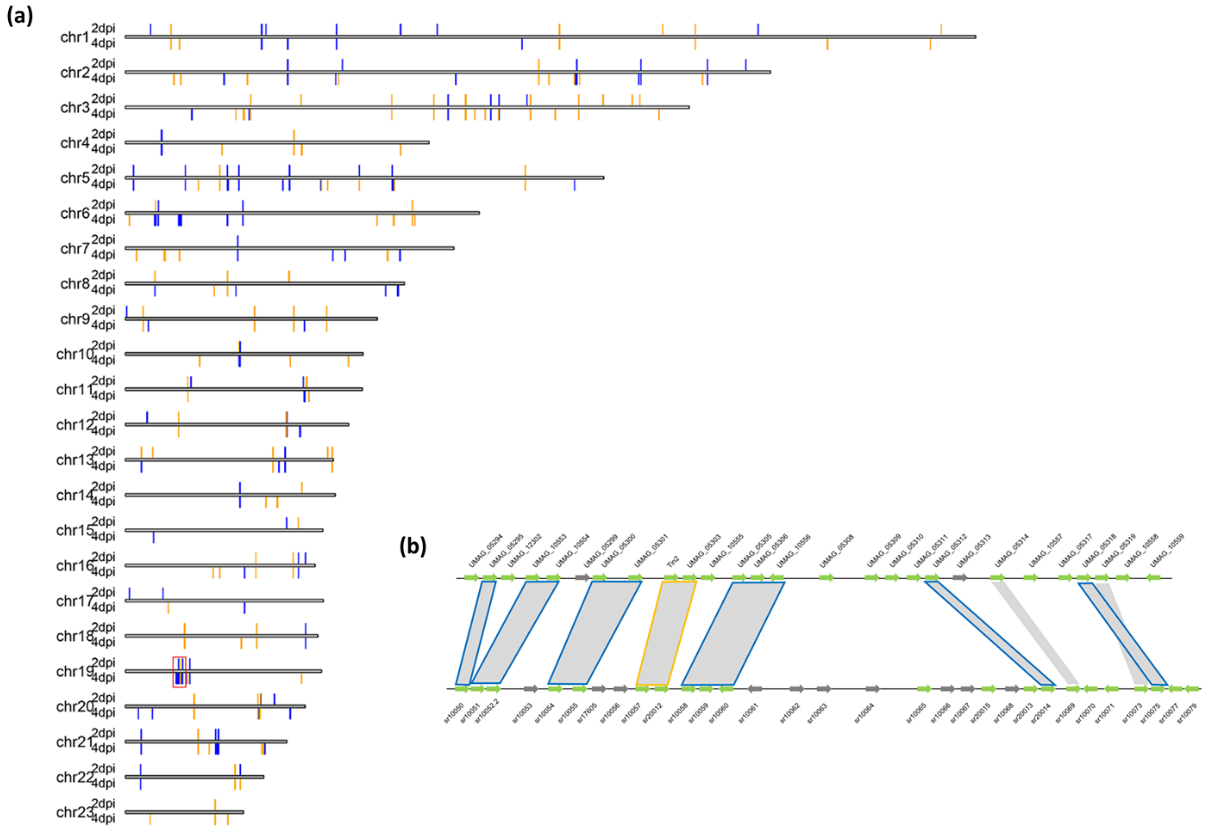


**Supplementary Fig.5 The distribution of effector DEOs on chromosomes**. **a,** the distribution of DEOs in 22 chromosomes based on the *U. maydis* genome organization. The blue bar shows *U. maydis* highly expressed DEOs, and the yellow bar shows the *S. reilianum* highly expressed DEOs. 2 dpi DEOs and 4 dpi DEOs were plotted above and below the chromosome, respectively. The red rectangle shows effector cluster 19A. **b,** the diagram shows the overview of synteny of effectors in cluster 19A. The effector genes are showed in green arrows; non-effector genes are shown in dark gray. The synteny of effector genes are in grey shade, the outline color of the shade indicates the DEO. The *U. maydis* highly expressed effector ortholog pairs are outlined in blue, the *S. reilianum* highly expressed genes are outlined in yellow.

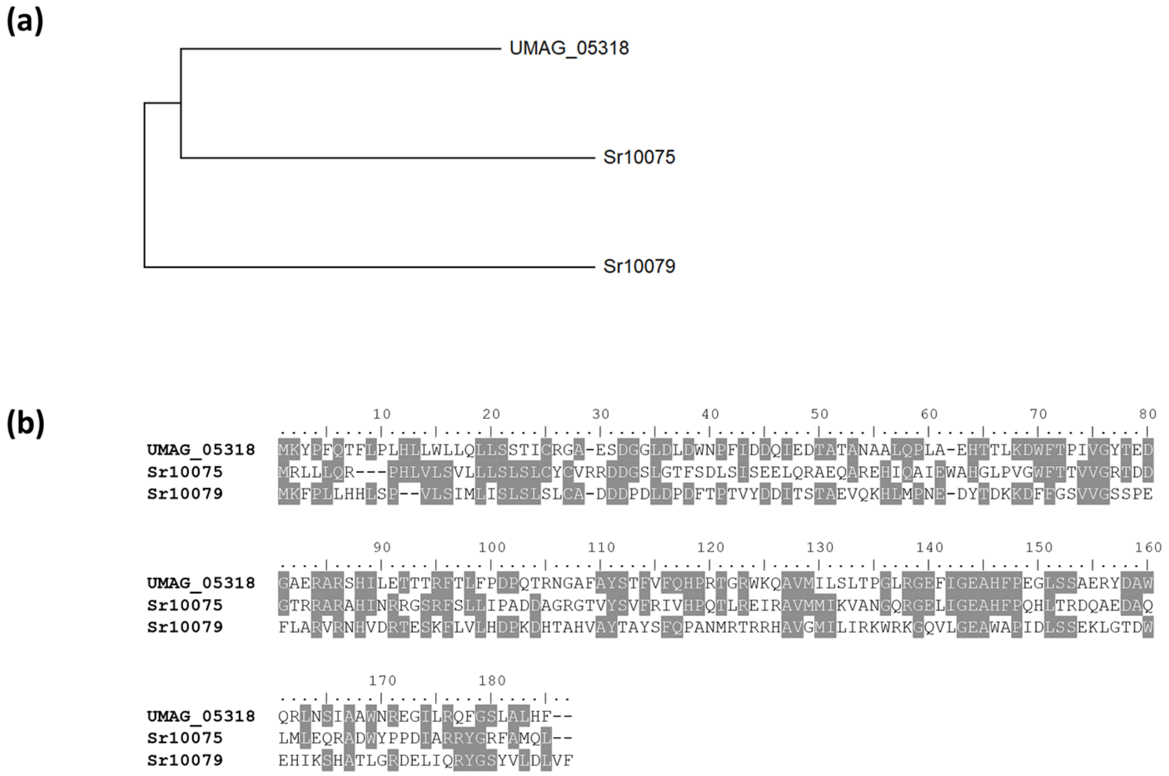


**Supplementary Fig.6 Sequence comparison of UMAG_05318, Sr10075 and Sr10079. a,** the maximum likelihood phylogenetic tree of UMAG_05318, Sr10075 and Sr10079 based on the amino acid sequences. Sr10079 is distant from UMAG_05318 and Sr10075. **b,** the amino acid sequence alignment of UMAG_05318, Sr10075 and Sr10079. The identical amino acids were shaded in grey.

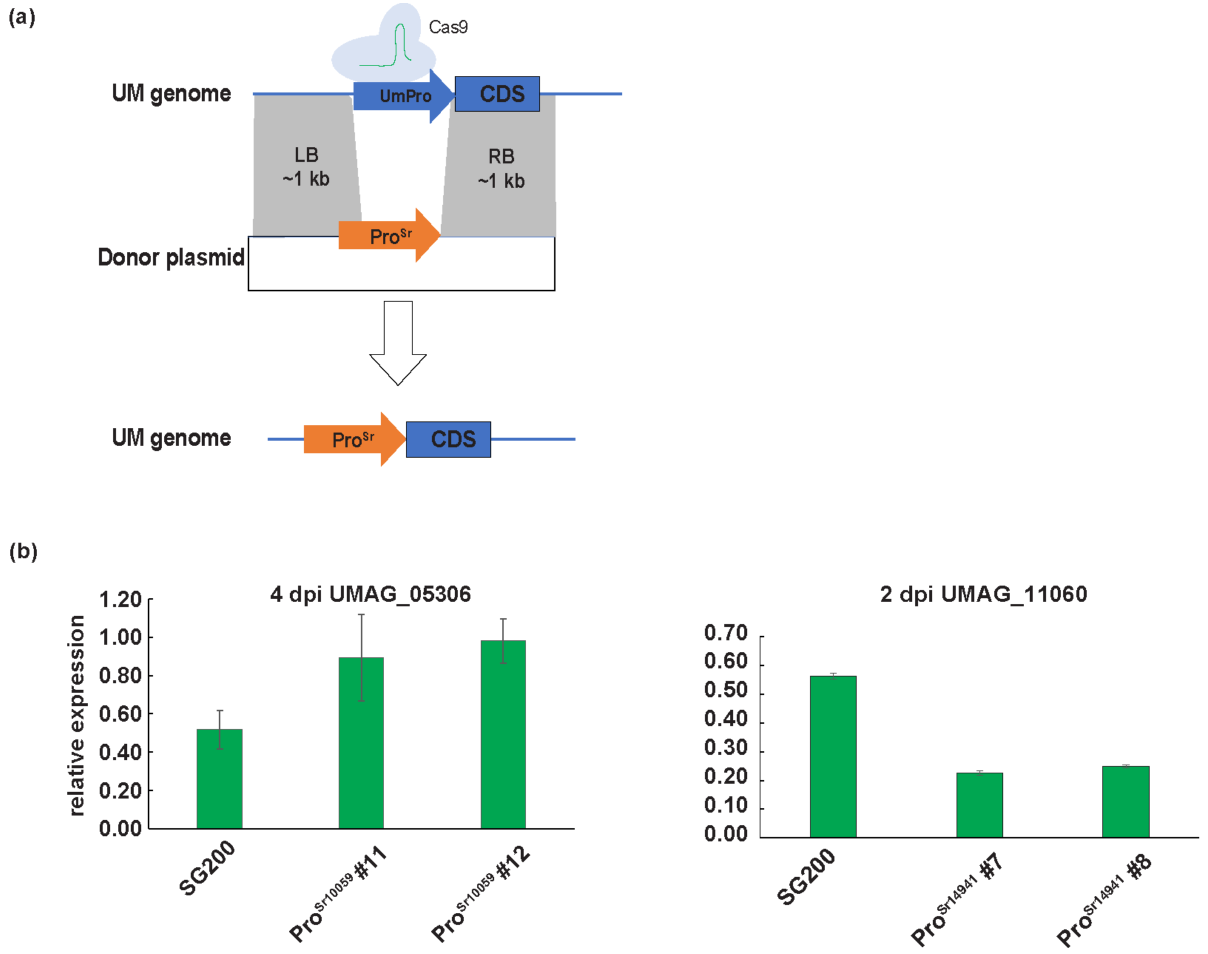


**Supplementary Fig. 7 The regulation of *S. reilianum* promoters in *U. maydis* by CRIPSR-Cas9 mediated promoter conversion. a,** Schematic model of the method used for CRISPR-Cas9 mediated promoter conversion. **b,** qPCR detection of the corresponding effectors expression under different orthologous *S. reilianum* promoters. At 4 dpi, Pro$^{sr10059}$ regulates gene expression to the similar level as Pro$^{UMAG_05306}$ in *U. maydis*, while at 2 dpi, Pro$^{sr14941}$ drove gene expression was significantly lower compared to Pro$^{UMAG_11060}$. Error bar indicates standard deviation from three biological replications.
